# Programmed downregulation of METTL3 is essential for decidualization in both humans and mice

**DOI:** 10.1101/2025.06.14.659666

**Authors:** Yu-Qi Hong, Run-Fu Jiang, Shi Tang, Xiao-Qi Yang, Meng Zhang, Yi Liu, Qing-Yan Zhang, Chen-Hui Ding, Zhan-Hong Zheng, Yan-Wen Xu, Shi-Hua Yang, Ji-Long Liu

## Abstract

*N*^6^-methyladenosine (m^6^A), the most abundant mRNA modification in eukaryotes, plays an essential role in regulating gene expression. Our prior research, alongside that of others, demonstrated that conditional uterine knockout of methyltransferase-like 3 (METTL3), the enzyme responsible for m^6^A modification, led to complete failure of embryo implantation and decidualization. Intriguingly, METTL3 expression is downregulated rather than upregulated in human endometrial stromal cells (HESCs) during in vitro decidualization and in mouse decidual tissues during pregnancy. We hypothesized that this decline in METTL3 expression is indispensable for successful decidualization. To test this hypothesis, we overexpressed METTL3 in HESCs and observed impaired decidualization in vitro. Additionally, we generated genetically engineered mice with uterine-specific METTL3 overexpression using *Pgr*-Cre, which exhibited subfertility mainly due to impaired decidualization. Further investigation revealed a marked decrease in HAND2, a well-established regulator of decidualization, following METTL3 overexpression. Mechanistically, we uncovered that METTL3 destabilizes *HAND2* mRNA via m^6^A modification at the 5′-UTR. In summary, our study underscores the critical role of programmed METTL3 downregulation in decidualization by sustaining HAND2 expression.

## Introduction

Decidualization is a critical prerequisite for embryo implantation and the maintenance of human pregnancy. Under the control of ovarian-derived 17β-estradiol (E_2_) and progesterone (P_4_), this process spontaneously commences during the secretory phase of the menstrual cycle, originating around the spiral arteries [1]. During decidualization, endometrial stromal cells (fibroblasts) undergo proliferation and differentiation, transforming into large, rounded decidual cells. These cells are notably marked by the secretion of prolactin (PRL) and insulin-like growth factor-binding protein 1 (IGFBP1) [2, 3]. The progression of decidualization correlates with the extent of trophoblast invasion, thus exerting a pivotal regulatory influence on embryo implantation, placental development, and the maintenance of pregnancy. Deficiencies in decidualization can precipitate various reproductive disorders, including unexplained infertility, recurrent miscarriage, intrauterine growth restriction, preeclampsia, and preterm birth [4]. A deeper understanding of the molecular mechanisms underlying decidualization may provide more clues and theoretical foundations for the prevention and treatment of infertility and pregnancy-related disorders.

Recently, *N*^6^-methyladenosine (m^6^A), the most abundant modification of mRNA in eukaryotes, has attracted widespread attention for its pivotal role in regulating gene expression [5]. By binding with different reader proteins, m^6^A modification regulates gene expression by modulating various mRNA processes, such as splicing, translocation, degradation, stabilization, and translation [6]. This modification is inherently dynamic and reversible, installed by a methyltransferase complex with methyltransferase-like 3 (METTL3) as its catalytic subunit [7], and removed by two demethylases: fat mass and obesity-associated protein (FTO) and alkB homolog 5 (ALKBH5) [8, 9]. Our previous research [10], alongside another study [11], has shown that loss of m^6^A modification in mouse uterus by conditional knockout of METTL3 with *Pgr*-Cre leads to a failure in both embryo implantation and decidualization. Intriguingly, we observed that METTL3 expression is downregulated, rather than upregulated, from the proliferative to the secretory phase in the human endometrium in vivo [10], as well as in cultured human endometrial stromal cells (HESCs) during in vitro decidualization [12]. Analysis of a public spatiotemporal transcriptomic dataset [13] revealed progressive METTL3 downregulation in mesometrial decidual cells or decidua basalis in the mouse uterus throughout pregnancy, from GD8 to GD18 (**Fig. S1**). These observations suggest that METTL3 downregulation occurs during decidualization in both humans and mice. However, the essentiality of this programmed METTL3 downregulation for decidualization remains to be definitively established.

Here, utilizing cultured human endometrial stromal cells (hESCs) and murine models, we found that forced METTL3 overexpression impaired decidualization. Further investigation revealed that METTL3 overexpression is accompanied by a marked decrease in HAND2 expression, a well-established regulator of decidualization. Mechanistic investigations elucidated that METTL3 overexpression destabilizes HAND2 mRNA via m^6^A modification at the 5′-UTR. Collectively, our study provides evidence that programmed downregulation of METTL3 is essential for decidualization.

## Results

### Overexpression of METTL3 impairs human endometrial stromal cell (HESC) decidualization in vitro

To investigate the effect of METTL3 overexpression on decidualization, we first utilized an in vitro system where primary human endometrial stromal cells (HESCs) were stimulated to differentiate using MPA and 8-Br-cAMP. METTL3 was overexpressed 6 hours prior to decidualization induction. Quantitative RT-PCR demonstrated a significant suppression of prolactin (PRL) and insulin-like growth factor binding protein 1 (IGFBP1), key markers of decidualization [4], following METTL3 overexpression after a 4-day decidualization period (**Fig. 1A**). Western blot analysis further confirmed the downregulation of IGFBP1 at the protein level (**Fig. 1B**). While HESCs typically assume a polygonal epithelioid morphology upon decidualization, METTL3 overexpression markedly impaired this transformation, as evidenced by reduced cell shape changes (**Fig. 1C**). Collectively, these results indicate that METTL3 overexpression disrupts decidualization in HESCs in vitro.

**Figure 1.**
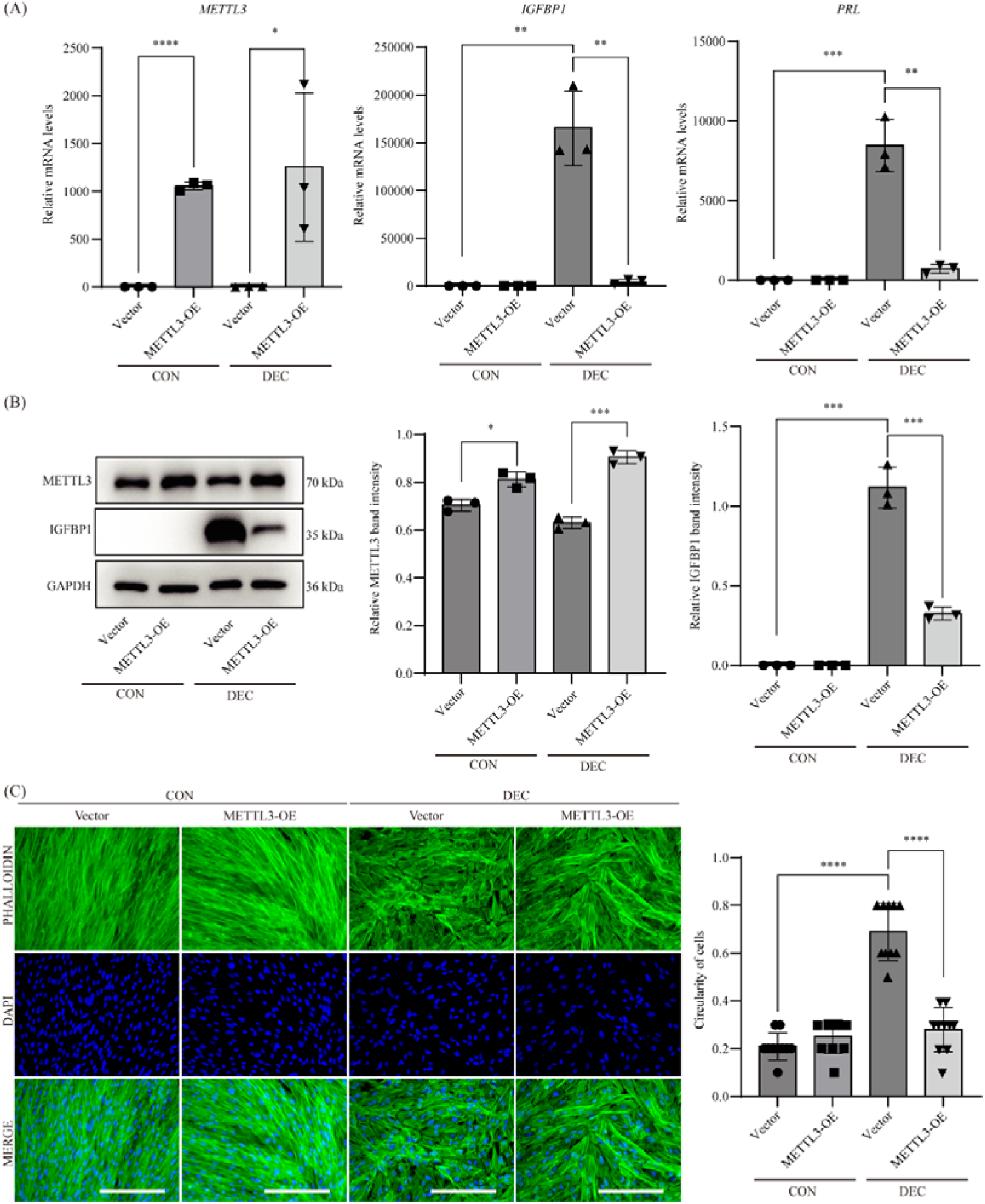
Over-expression of METTL3 impairs human endometrial stromal cell (HESC) decidualization in vitro. (A) Quantitative RT-PCR analysis of PRL and IGFBP1 expression in primary HESCs following METTL3 overexpression in the in vitro decidualization model. CON, vehicle control; DEC, in vitro decidualization for 4 days. Data are presented as means ± SEM. *P < 0.05; **P < 0.01; ***P < 0.001; ****P < 0.0001. (B) IGFBP1 expression in primary HESCs following METTL3 overexpression in the in vitro decidualization model. Data are presented as means ± SEM. *P < 0.05; **P < 0.01; ***P < 0.001. (C) Phalloidin staining of F-actin in primary HESCs following METTL3 overexpression in the in vitro decidualization model. Circularity = (4 * π * Area) / (Perimeter^2), where Area is the area of the cell and Perimeter is the length of the cell’s boundary. The resulting value ranges from 0 to 1, where 1 represents a perfect circle and lower values indicate increasingly irregular shapes. Bar = 100 μm. Data are presented as means ± SEM. ****P < 0.0001.

### Mice with uterine-specific METTL3 overexpression show subfertility

In order to further understand the role of m^6^A modification in the uterus, we generated mice with METTL3 overexpression in the uterus (METTL3-OE) by crossing CAG-LSL-Mettl3 mice with *Pgr*-Cre mice (**Fig. 2A**). Littermate CAG-LSL-Mettl3 mice served as wild-type control (**Fig. 2B**). PCR analysis confirmed Cre-mediated recombination of loxp sites in METTL3-OE mouse uterus (**Fig. S2A-C**). The overexpression of METTL3 in the uterus of METTL3-OE mice was further confirmed by quantitative RT-PCR (**Fig. 2C**), western blot (**Fig. 2D**) and immunohistochemistry (**Fig. 2E**). Moreover, we performed m^6^A dot blot analysis and found that m^6^A methylation was significantly increased in the uterus of METTL3-OE mice (**Fig. 2F**). These results verified the success of METTL3 overexpression in the uterus of METTL3-OE mice.

**Figure 2.**
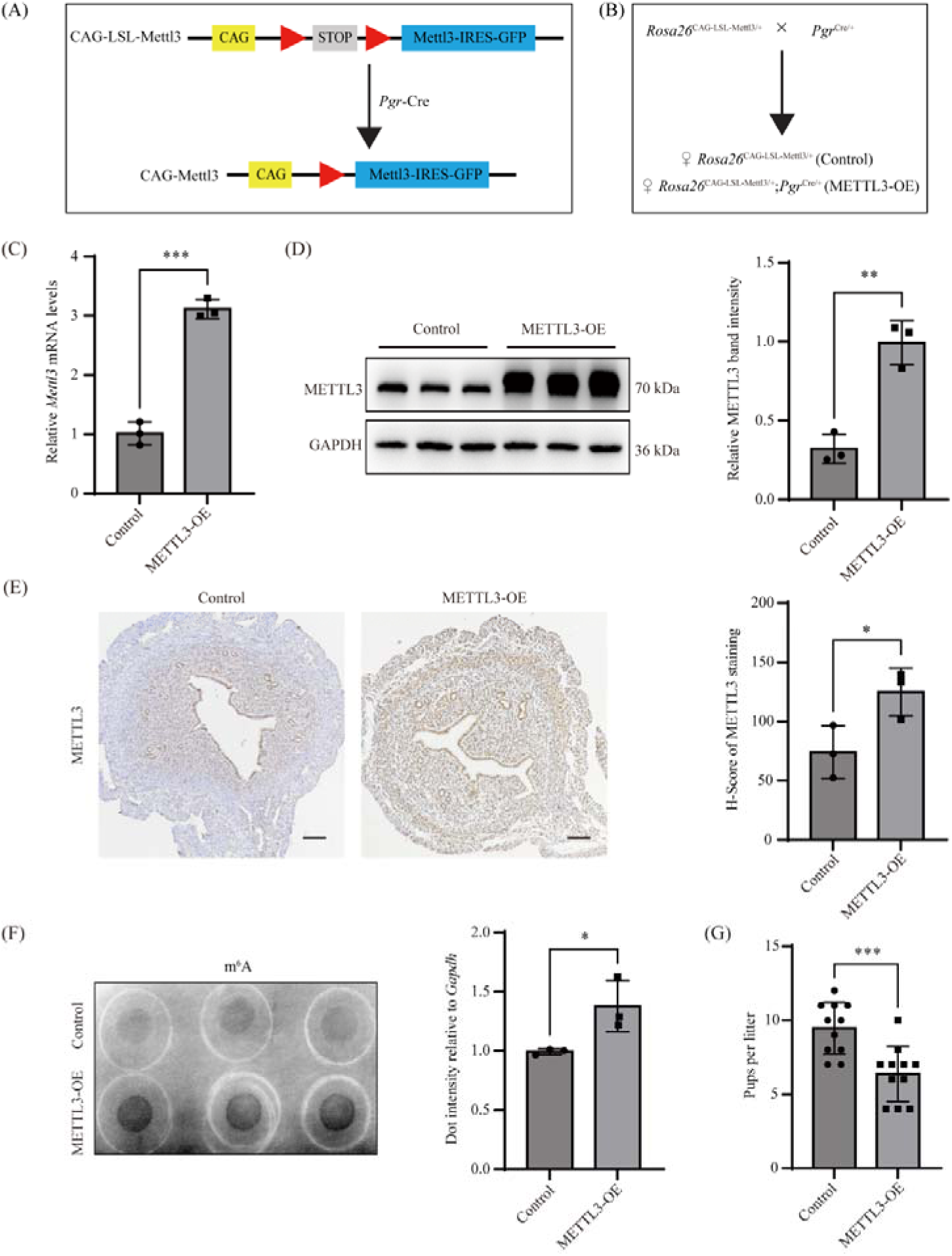
***Pgr*-Cre-mediated METTL3 overexpression in mice leads to subfertility.** (A) Diagram showing the strategy to generate METTL3 overexpression allele at the *Rosa26* locus. (B) A diagram of the breeding scheme. Uterine-specific METTL3 overexpression (METTL3- OE) mice were generated by crossing *Rosa26*^CAG-LSL-Mettl3/+^ mice with the *Pgr*-Cre driver line. (C) Quantitative RT-PCR analysis of *Mettl3* mRNA levels in the uterus of METTL3-OE and control mice on GD4. Data are presented as mean ± SEM. ***P < 0.001. (D) Western blot analysis of METTL3 protein levels in the uterus of METTL3-OE and control mice on GD4. Data are presented as mean ± SEM. **P < 0.01. (E) Immunohistochemical staining of METTL3 protein in the uterus of METTL3-OE and control mice on GD4. Bar = 100 μm. Data are presented as mean ± SEM. *P < 0.05. (F) Dot blot showing the m^6^A levels of mRNA in the uterus of METTL3-OE and control mice on GD4. Dot intensity was normalized to qRT-PCR-determined *Gapdh* levels. Data are presented as mean ± SEM. *P < 0.05. (G) Bar plot showing the 6-mo fertility test results for METTL3-OE and control mice. Data are presented as mean ± SEM. ***P < 0.001.

We then conducted a fertility test to evaluate the effect of METTL3 overexpression in mice. METTL3-OE and control females were mated with wild-type males of proven fertility. During a 6-month fertility test, both METTL3-OE and control females had normal reproductive cycles and exhibited normal mating behavior with the formation of vaginal plugs. However, METTL3-OE females produced significantly fewer pups per litter than control females (9.455 ± 0.529 pups/litter v.s. 6.364 ± 0.560 pups/litter; mean ± SEM; P=0.0007) (**Fig. 2G**).

### Overexpression of METTL3 leads to impaired decidualization

To determine the cause of subfertility in METTL3-OE mice, we first assessed ovarian function. Histological examination of ovaries on gestational day 4 (GD4) showed no overt morphological differences between METTL3-OE and control mice, both showing the presence of corpus lutea (**Fig. 3A**). Serum E_2_ and P_4_ levels were also comparable between METTL3-OE and control mice (**Fig. 3B**). Additionally, immunohistochemistry analysis showed that the expression of key steroid biosynthetic enzymes, cytochrome P450 cholesterol side-chain cleavage enzyme (P450scc/CYP11A1) and 3β-hydroxysteroid dehydrogenase (3β- HSD/HSD3B2) [14], were unaffected in the developing corpus luteum of METTL3-OE mice (**Fig. 3C**). These results indicate that ovary function was normal in METTL3-OE mice.

**Figure 3.**
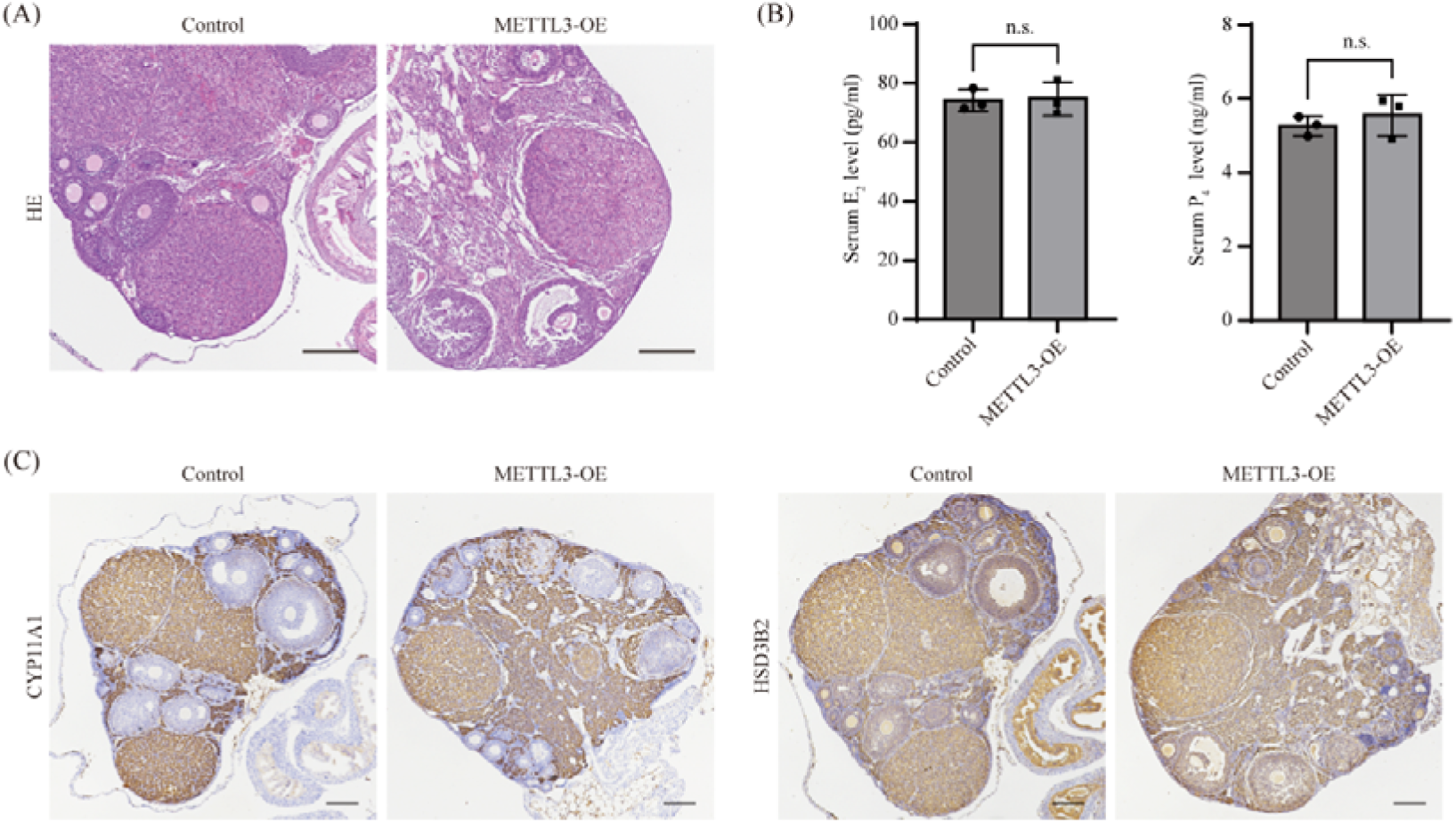
Post-ovulation corpus luteum development is unaffected in METTL3-OE mice. (A) HE staining of the ovary from METTL3-OE mice on gestational day 4 (GD4). Bar = 100 μm. (B) The concentration of circulating E_2_ and P_4_ in METTL3-OE mice on GD4. Data are presented as mean ± SEM. n.s., not significant. (C) Immunohistochemical staining of CYP11A1 and HSD3B2 in the ovary of METTL3-OE mice on gestational day 4 (GD4). Bar = 100 μm.

We then surveyed the pregnancy status of these mice at different time points. On GD4, the morphology of the uterus appeared indistinguishable between METTL3-OE and control mice (**Fig. 4A**). Embryo implantation occurred normally on GD5 in METTL3-OE mice and the number of implantation sites was comparable between METTL3-OE and control mice (**Fig. 4B**). On GD8, although the number implantation sites were also comparable, the weight of implantation sites was significantly reduced in METTL3-OE mice compared with control mice (**Fig. 4C**). Alkaline phosphatase activity, a marker of uterine stromal cell decidualization [3], was downregulated at the anti-mesometrial region of implantation sites in METTL3-OE uteri (**Fig. 4D**). In addition, decidualization defect in METTL3-OE mice was validated by the downregulation of two decidualization marker genes, prolactin family 8 subfamily A member 2 (*Prl8a2*) [15] and bone morphogenetic protein 2 (*Bmp2*) [16]. Notably, the expression of another decidualization marker gene, Wnt family member 4 (*Wnt4*) [17], was unchanged (**Fig. 4E**).

**Figure 4.**
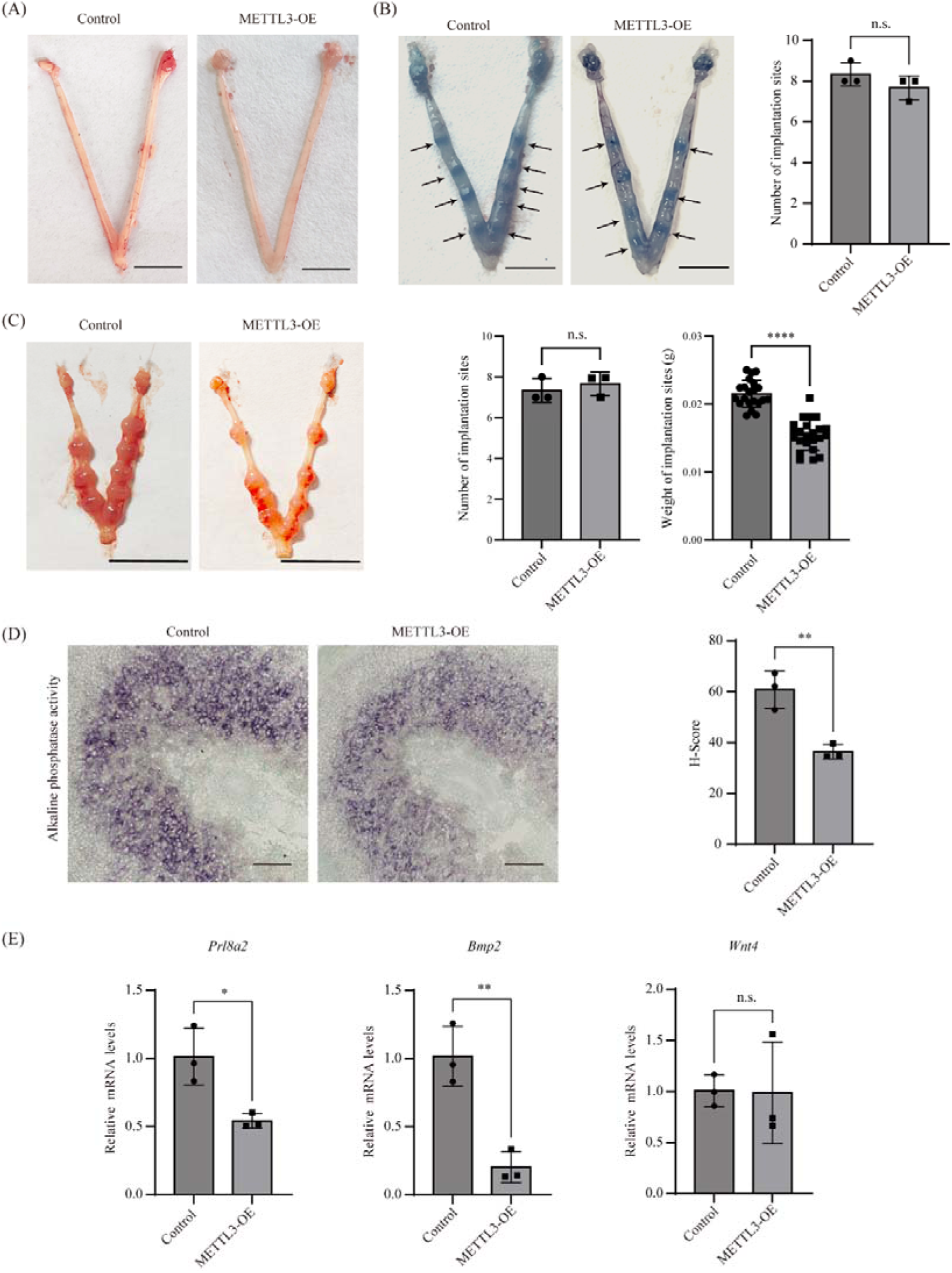
METTL3 showed normal embryo implantation, but impaired decidualization. (A) Photos of the uterus from the uterus of METTL3-OE mice on GD4. Bar = 1 cm. (B) Bar plot showing the number of embryo implantation sites in METTL3-OE mice on GD5. Data are presented as mean ± SEM. n.s., not significant. Representative photos of the uterus are shown on the left panel. Bar = 1 cm. Implantation sites are marked by arrowheads. (C) Bar plots showing number of implantation sites per mouse and weight of implantation sites in METTL3-OE mice on GD8. Bar = 1 cm. Data are presented as mean ± SEM. ***P < 0.001. (D) Alkaline phosphatase staining of the implantation site in in METTL3-OE mice on GD. Bar = 100 μm. Data are presented as mean ± SEM. **P < 0.01. (E) Quantitative RT-PCR analysis of *Prl8a2*, *Bmp2* and *Wnt4* at the implantation site of METTL3-OE mice on GD8. Data are presented as mean ± SEM. *P < 0.05; **P < 0.01. n.s., not significant.

To further confirm the decidualization defect in METTL3-OE mice, we employed an artificial decidualization model. Sesame seed oil was injected into the lumen of one uterine horn to trigger decidual response instead of embryo implantation, with the contralateral horn serving as the unstimulated control. Based on the weight ratio of uterine horns, decidual response was significantly reduced in METTL3-OE mice (**Fig. S3A**). Decidualization defect in this model was verified by the downregulation of alkaline phosphatase activity (**Fig. S3B**) and mRNA markers *Prl8a2* and *Bmp2* (**Fig. S3C**).

### METTL3 overexpression results in increased PGR expression and decreased HAND2 expression

To unravel the molecular mechanisms underlying the compromised decidualization observed in METTL3-OE mice, we conducted a thorough investigation focusing on two critical timepoints in the uterus: GD4, prior to decidualization, and GD8, subsequent to decidualization.

We examined a panel of protein markers in METTL3-OE uteri on GD4, including steroid receptors (ESR1 and PGR), receptivity markers (MKI67 and MUC1), epithelial cell markers (TACSTD2 and FOXA2) and stromal cell markers (HOXA11 and HAND2) [18–20]. Previously, we found that *Pgr*-Cre mediated deletion of *Mettl3* in the uterus results in decreased expression of PGR at the protein level [10]. Consistently, we here discovered that although ESR1 protein was unchanged (**Fig. 5A**), PGR protein was significantly up-regulated in METTL3-OE uteri (**Fig. 5B**). Immunohistochemistry staining for MKI67 demonstrated that epithelial cell proliferation had ceased in both METTL3-OE and control uteri (**Fig. 5C**), indicating normal uterine receptivity. This finding was further corroborated by comparable levels of MUC1 in the uterine epithelial cells of METTL3-OE mice (**Fig. 5D**). Nevertheless, a noteworthy observation was the significantly reduced number of proliferative stromal cells in the METTL3-OE uteri (**Fig. 5C**). This decrease may provide a clue to the subsequent impairment in decidualization observed in these mice. The epithelial cell structure appeared normal in the uteri of METTL3-OE mice, as evidenced by the expression patterns of luminal cell marker TACSTD2 (**Fig. 5E**) and glandular cell marker FOXA2 (**Fig. 5F**). Interestingly, the expression of TACSTD2 was significantly lower in METTL3-OE uteri than control uteri. Given that TACSTD2 is negatively regulated by P_4_ and PGR in the mouse uterus [10, 21], this finding corroborates the PGR overexpression phenotype observed in METTL3-OE uteri (**Fig. 5B**). HOXA11 serves as a pan-stromal cell marker, while HAND2 specifically identifies superficial stromal cells. In our analysis, we observed that while the expression of HOXA11 was unchanged (**Fig. 5G**), the expression of HAND2 was significantly downregulated in the uteri of METTL3-OE mice (**Fig. 5H**). This downregulation of HAND2 was also supported by quantitative RT-PCR (**Fig. S3A**) and western blot (**Fig. S3B**). Given the crucial role of HAND2 in mouse implantation and decidualization [22, 23], this downregulation of HAND2 may represent an important defect in the METTL3-OE uteri. Intriguingly, despite this downregulation, uterine receptivity and subsequent implantation appeared to be unaffected in METTL3-OE uteri.

**Figure 5.**
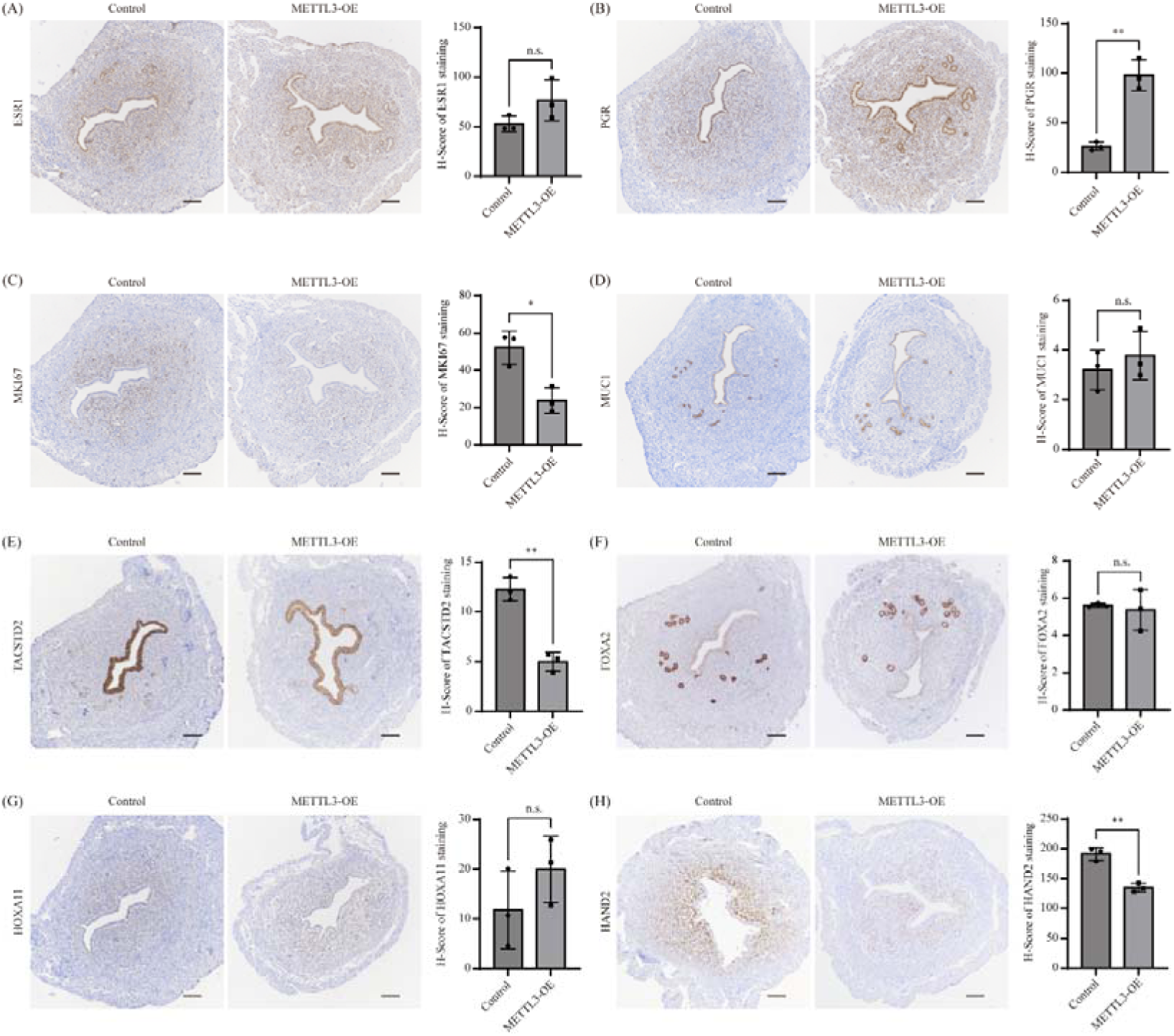
Overexpression of METTL3 leads to increased PGR expression and decreased HAND2 expression on GD4. Shown are immunohistochemical staining of ESR1 (A), PGR (B), MKI67 (C), MUC1 (D), TACSTD2 (E), FOXA2 (F), HOXA11 (G) and HAND2 (H) in the uterus of METTL3-OE mice on GD4. Bar = 100 μm. Data are presented as mean ± SEM. *P < 0.05; **P < 0.01.

To explore the causes of impaired decidualization in METTL3-OE mice, we collected uterine tissues specifically from the implantation site on GD8. The fetal tissues were discarded under a stereo microscope as described previously [24]. RNA-seq analysis showed that 342 genes were upregulated and 226 genes were downregulated in METTL3-OE uteri in comparison to control uteri (**Fig. 6A** and **Table S1**). According to gene ontology analysis, cell differentiation was significantly enriched among downregulated genes (**Fig. 6B**). Previously, we obtained a list of known decidualization-related genes from the PubMed database by using a text mining approach [25]. By comparing our RNA-seq data against this list, we found that the expression of HAND2 was downregulated (**Fig. 6C**). This finding was further confirmed by quantitative RT-PCR (**Fig. 6D**), western blot (**Fig. 6E**) and immunohistochemistry (**Fig. 6F**). These results suggest that the reduced expression of HAND2 in decidual cells might be a contributing factor to the observed decidualization impairment in METTL3-OE mice.

**Figure 6.**
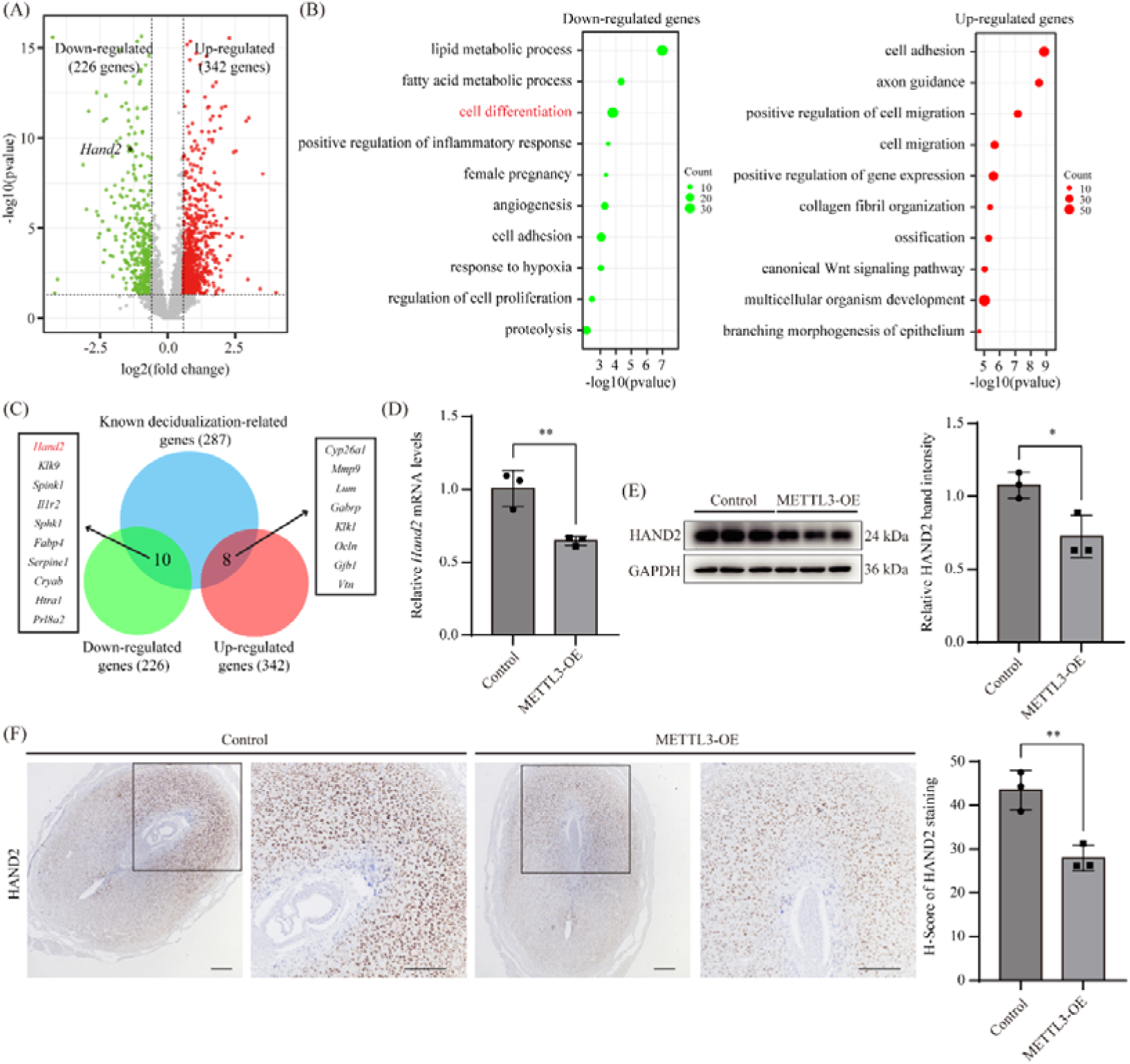
METTL3 overexpression results in reduced HAND2 expression at the implantation sites of mouse uterus on GD8. (A) Volcano plot showing significantly expressed genes at the implantation sites of mouse uterus on GD8 in METTL3-OE mice. Differentially expressed genes were selected based on the criteria of fold change > 2 and P < 0.05. (B) Gene ontology enrichment analysis of down-regulated genes and up-regulated genes, respectively. (C) Venn diagram depicting the overlap of differentially expressed genes and known genes associated with mouse decidualization. (D) Quantitative RT-PCR analysis of *Hand2* mRNA levels at the implantation sites of mouse uterus on GD8 in METTL3-OE mice. Data are presented as mean ± SEM. **P < 0.01. (E) Western blot analysis of HAND2 protein levels at the implantation sites of mouse uterus on GD8 in METTL3-OE mice. Data are presented as mean ± SEM. *P < 0.05. (F) Immunohistochemical staining of HAND2 at the implantation sites of mouse uterus on GD8 in METTL3-OE mice. Bar = 100 μm. Data are presented as mean ± SEM. **P < 0.01.

### METTL3 negatively regulates HAND2 expression by m^6^A modification at the 5**′**-UTR

METTL3, as the core subunit of the m^6^A methyltransferase complex, primarily facilitates the formation of m^6^A modifications on mammalian mRNAs [7]. The m^6^A modification has been reported to regulate many aspects of mRNA biology; however, its major function is to promote mRNA degradation in the cytoplasm [26]. Since HAND2 is downregulated by METTL3 in mouse uterus, we examined if mouse *Hand2* mRNA is a direct substrate for m^6^A modification. By data mining of our previous MeRIP-seq dataset on mouse uterus (GSE211521), we identified two m^6^A peaks spanning almost the entire *Hand2* mRNA (**Fig. 7A**). Similar results were obtained by data mining of a MeRIP-seq dataset performed on human endometrium (GSE205398) (**Fig. 7B**). Furthermore, we found that, in primary HESCs, expression of HAND2 mRNA was significantly inhibited when METTL3 was overexpressed (**Fig. 7C**), which was consistent with the findings from mouse uteri (**Fig. 5H** and **Fig. 6F**). These results indicated that the METTL3-HAND2 axis is likely conserved between mice and humans.

**Figure 7.**
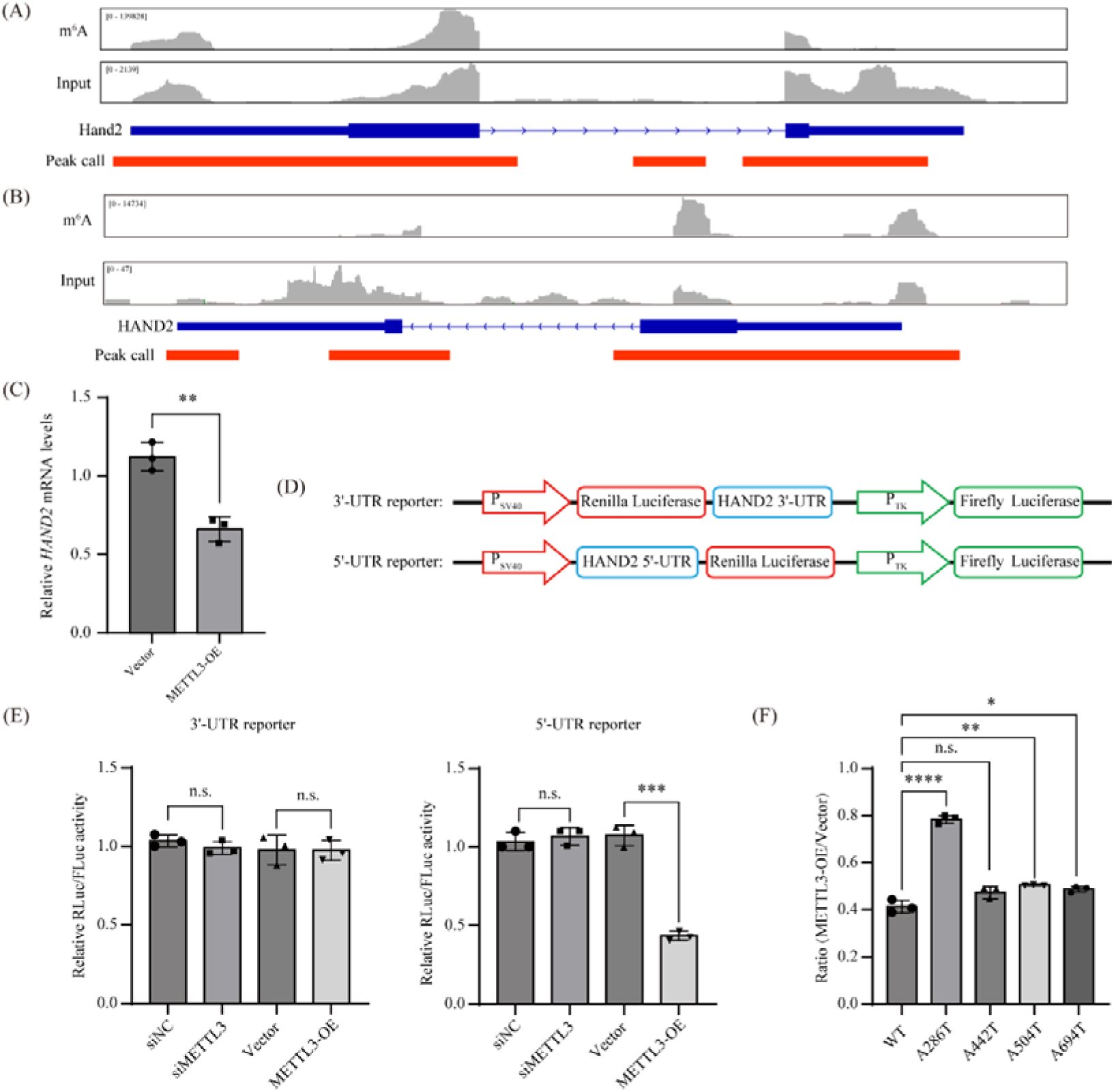
**METTL3 negatively regulates HAND2 expression via m^6^A modification of *HAND2* 5**′**-UTR at A286.** (A) Integrative genomics viewer (IGV) illustrating the m^6^A peak distribution along the mouse Hand2 mRNA in MeRIP-seq dataset GSE205398. Peak calling was performed by using the MACS3 software. (B) Integrative genomics viewer (IGV) illustrating the m^6^A peak distribution along the human HAND2 mRNA in MeRIP-seq dataset GSE205398. Peak calling was performed by using the MACS3 software. (C) Quantitative RT-PCR analysis showing that HAND2 mRNA expression is down-regulated in HESCs following METTL3 overexpression. (D) Schematic diagram of dual luciferase assay using the psiCHECK^TM^-2 vector. The 3′-UTR reporter is constructed by inserting HAND2 3′-UTR sequence downstream of the Renilla luciferase (RLuc) gene. The 5′-UTR reporter is constructed by inserting HAND2 5′-UTR sequence between the SV40 promoter and the RLuc gene. The firefly luciferase gene (FLuc), driven by the TK promoter, served as a normalization control for transfection efficiency. (E) The relative RLuc/FLuc activities in HEK293T cells co-transfected with the dual luciferase reporter and either METTL3-OE or vector control for 48 h. Data are presented as mean ± SEM. ***P < 0.001. (F) The impact of A-to-T mutation at potential m^6^A sites in the 5′-UTR reporter. Data are presented as mean ± SEM. *P < 0.05; **P < 0.01; ****P < 0.0001.

Utilizing the SRAMP tool [27], we predicted the existence of five potential m^6^A modification sites (A286, A1274, A1663, A1697, and A2394) within the human *HAND2* mRNA (**Fig. S5A**). Notably, three of these predicted sites (A286, A1663, and A1697) exhibited conservation between mice and humans (**Fig. S5B**). Specifically, A286 is located within the 5′-UTR, whereas A1663 and A1697 are situated in the 3′-UTR. To further investigate the impact of these conserved sites, we constructed a 5′-UTR reporter and a 3′-UTR reporter (**Fig. 7D**). The 5′-UTR reporter is constructed by inserting HAND2 5′-UTR sequence between the SV40 promoter and the Renilla luciferase (RLuc) gene. The 3′-UTR reporter is constructed by inserting HAND2 3′-UTR sequence downstream of the RLuc gene. The firefly luciferase gene (FLuc), driven by the TK promoter, served as a normalization control for transfection efficiency. The luciferase assay demonstrated that the 3′-UTR reporter did not respond to either siMETTL3 or METTL3 overexpression. Interestingly, despite the fact that the relative RLuc/FLuc activity remained unchanged in the 3′-UTR reporter upon METTL3 knockdown, it was significantly reduced following METTL3 overexpression (**Fig. 7E**). In order to identify the precise locations of m^6^A modification within the 5′-UTR, we generated A-to-T point mutations in the 5′-UTR reporter to abolish each potential m^6^A modification sites. Besides the A286 site predicted by the SRAMP tool, we also tested another three potential sites (A442, A504 or A694) by searching the entire HAND2 5′-UTR for the GGAC motif [10]. The A286T mutation, but not A442T, A504T or A694T, could effectively change the relative RLuc/FLuc activity compared with wild-type control (**Fig. 7F**). Our data suggest that METTL3 overexpression represses HAND2 by m^6^A modification at the A286 site in the 5′-UTR.

## Discussion

Previously, through studies employing uterine-specific knockout mice, our group and others have demonstrated that METTL3 is indispensable for embryo implantation and decidualization [10, 11]. Intriguingly, METTL3 expression decreases rather than increases during the decidualization process [10, 12]. Here, we reveal that this downregulation is also pivotal for decidualization, as evidenced by the finding that METTL3 overexpression disrupts decidualization. These findings underscore the complex role of METTL3 in regulating decidualization.

In this study, we employed both in vitro and in vivo models to investigate the role of METTL3 downregulation in decidualization. In human endometrial stromal cells (hESCs), we found that overexpression of METTL3 impaired decidualization. This finding is in line with our previous study showing that METTL3 knockdown before decidualization initiation (day 0) significantly impaired differentiation capacity, whereas METTL3 knockdown after decidualization initiation (day 2) unexpectedly had no effect on this process [12]. Given that METTL3 expression is downregulated during the decidualization process, our findings suggest that METTL3 expression in hESCs is essential for the initiation phase of decidualization, while excessive METTL3 expression in the late stage of decidualization is harmful. In *Pgr*-Cre-mediated METTL3 conditional overexpression mice, we observed that these mice were subfertile. The major phenotype of these mice was compromised decidualization. Conditional knockout of METTL3 in mouse uterus mediated by *Pgr*-Cre (*Mettl3*^d/d^) resulted in infertility. A complete failure in implantation and decidualization was observed in *Mettl3*^d/d^ mice [10, 11]. In METTL3-OE mice, implantation was normal, and while decidualization was impaired, it was not completely abolished. Therefore, it appears that the fertility phenotype observed in METTL3-OE mice is significantly less severe compared to that in *Mettl3*^d/d^ mice. There are two plausible explanations for this milder phenotype in METTL3-OE mice. Firstly, the expression of METTL3 in the uteri of METTL3- OE mice is only ∼threefold higher compared to control mice, indicating a relatively modest increase. Secondly, in METTL3-OE uteri, the eraser *Fto* is significantly up-regulated, while the readers *Ythdc2* and *Ythdf1* are down-regulated (as shown in **Fig. S6**). This suggests a feedback mechanism that limits the impact of METTL3 overexpression, thereby mitigating the severity of the fertility phenotype. Taken together, our findings suggest that METTL3 downregulation is pivotal for decidualization.

In this study, we pinpointed reduced HAND2 as a pivotal downstream effector of METTL3 overexpression. Specifically, we observed a targeted downregulation of HAND2 expression in METTL3-overexpressing (METTL3-OE) uteri. Given HAND2’s essential role in implantation and decidualization [22, 23], this decline likely signifies a functional impairment in METTL3-OE uteri. Mechanistically, we unveiled that METTL3 directly represses HAND2 expression by introducing m^6^A modifications at the 5′-UTR of *HAND2* mRNA. This discovery is particularly intriguing in light of our prior research, which demonstrated that METTL3 indirectly elevates HAND2 expression via PGR [10]. In that context, METTL3 modifies the 5′-UTR of *PGR* mRNA with m^6^A, boosting PGR protein translation, which subsequently promotes HAND2 transcription. In *Mettl3*^d/d^ uteri, the lack of METTL3 results in diminished PGR protein levels, thereby suppressing HAND2 transcription. Consequently, the scarcity of HAND2 mRNA renders the impact of METTL3-mediated m^6^A modification on HAND2 mRNA negligible. Conversely, in METTL3-OE uteri, the direct suppressive effect of METTL3 on HAND2 mRNA through m^6^A modification prevails. The influence of the METTL3-PGR-HAND2 axis is eclipsed by the direct action of METTL3 on HAND2 mRNA. Our findings are schematically illustrated in **Fig. 8**, offering a comprehensive overview of the intricate interplay between METTL3, PGR, and HAND2 in governing uterine function.

**Figure 8.**
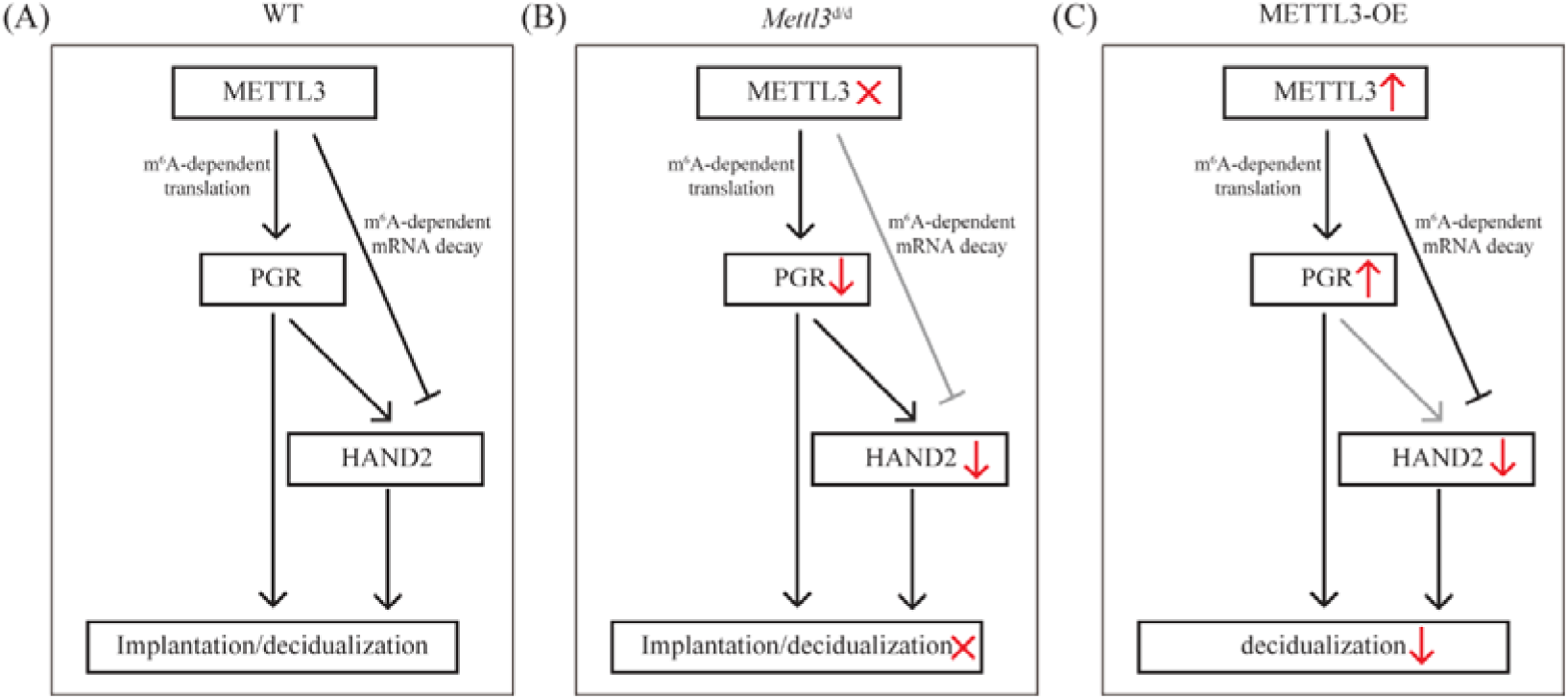
A proposed model. (A) In wild-type (WT) mice, METTL3 intricately regulates HAND2 expression through direct and indirect mechanisms. Directly, METTL3 reduces HAND2 expression by introducing m^6^A modification in the 5′-UTR of HAND2 mRNA, thereby accelerating its mRNA decay. Indirectly, METTL3 enhances HAND2 transcription via PGR. This is achieved by installing m^6^A modification in the 5′-UTR of PGR mRNA, which boosts PGR protein translation. PGR in turn enhances HAND2 transcription. (B) In mice with uterine-specific deletion of METTL3 (*Mettl3*^d/d^), the absence of METTL3 leads to a decrease of PGR protein expression, which subsequently curtails HAND2 transcription. Consequently, the scarcity of HAND2 mRNA as a substrate renders the impact of METTL3- mediated m^6^A modification on HAND2 mRNA negligible. (C) Conversely, in mice with METTL3 overexpression in the uterus (METTL3-OE), METTL3 directly suppresses HAND2 mRNA levels through m^6^A modification. In this scenario, the influence of the METTL3-PGR- HAND2 axis is overwhelmed by the direct action of METTL3 on HAND2 mRNA.

Remarkably, our findings may hold substantial implications for human reproductive health. A microarray dataset (GSE111974) revealed an elevation of METTL3 in the endometrium of patients experiencing recurrent implantation failure (RIF) during the implantation window [28]. This observation was strengthened by another study, which employed quantitative RT-PCR to detect an upregulation of METTL3 in RIF patients [29]. However, intriguingly, a microarray dataset (GSE58144) [30] referenced in a prior study [11] pointed to a downregulation of METTL3 in RIF patients. Downregulation of METTL3 in RIF patients was also supported by an RNA-seq dataset (GSE207362) [31] as we described previously [12]. The rationale behind this apparent discrepancy remains unclear. A plausible explanation is that METTL3 expression may exhibit heterogeneity among RIF patients, ranging from high to low levels. Given that both elevated and diminished METTL3 expressions can impair uterine function and female fertility, our hypothesis warrants further investigation.

In summary, our study demonstrates that METTL3 downregulation is critical for decidualization, as evidenced by the disruption of this process following METTL3 overexpression, both in vitro using human endometrial stromal cells and in vivo within the mouse uterus. These findings provide valuable insights into the mechanisms governing decidualization by elucidating the intricate relationship between gene expression patterns and gene function during this critical reproductive process.

## Materials and methods

### In vitro decidualization model using primary human endometrial stromal cells (HESCs)

Three normal fertile participants who had no apparent endometrial pathology and had a confirmed clinical pregnancy after embryo transfer were recruited for endometrial biopsy in the First Affiliated Hospital of Sun Yat-sen University in China from July 2020 to October 2021. This study had been approved by the Ethical Committee of the First Affiliated Hospital of Sun Yat-sen University (No. 2018-266) and all participants signed an informed consent. Detailed information of patients is listed in **Table S2**. The endometrial tissues were cut into the smallest possible pieces and then subjected to digestion with type I collagenase (Gibco) for a duration of 1 hour. The epithelial and stromal cells were carefully separated using membrane filters with pore sizes of 100 µm and 40 µm (Corning). Subsequently, the human endometrial stromal cells (HESCs) were cultured in DMEM/F12 medium (Gibco) supplemented with 10% charcoal-stripped fetal bovine serum (cFBS, VivaCell). To induce the process of decidualization, the cells were treated with a combination of 0.5 mM 8-Br-cAMP (Sigma) and 1 µM medroxyprogesterone acetate in 2% cFBS for a period of 4 days.

### Mice

*Rosa26*^CAG-LSL-Mettl3/+^ mice (Cat. No. NM-KI-190056) and *Pgr*^Cre/+^ mice (Cat. No. NM-KI- 200117) were purchased from Shanghai Model Organisms Center, Inc. To generate uterine-specific METTL3 overexpression (METTL3-OE) mice, we crossed the *Rosa26*^CAG-LSL-Mettl3/+^ mice with the *Pgr*^Cre/+^ mice. The *Rosa26*^CAG-LSL-Mettl3/+^ littermates served as the wild-type control for this study. All mice were bred under specific pathogen-free conditions with a 12-h day and 12-h night cycle. All the animal procedures were approved by the Institutional Animal Care and Use Committee of South China Agricultural University (No. 2021B036).

### Fertility test

METTL3-OE and control mice were cohoused in cages with fertile wild-type males, maintaining a 1:1 ratio. Over a 6-month breeding period, the animals were carefully monitored every three days. Promptly after weaning, the pups were separated, and a meticulous record was kept of the number of pups per litter as well as the total number of litters born.

### Timed mating experiments

Female METTL3-OE and control mice were paired overnight with fertile wild-type males in separate cages. The day of the presence of a vaginal plug was designated as gestational day 1 (GD1). Once plug-positive, the females were separated from the males until the designated time for experimentation. To confirm pregnancy on GD4, one uterine horn of each plug-positive female was gently flushed with saline to verify the presence of blastocysts. If blastocysts were detected, the contralateral horn was selected for further experiments; otherwise, the mice were excluded from the study. On GD5, implantation sites in the uterus were visualized by intravenously injecting a blue dye solution (1% Chicago Blue in Saline, 100 µL/mouse), followed by a 4-minute wait before sacrificing the mice.

### Artificial decidualization

Pseudo-pregnancy was induced in female METTL3-OE and control mice through mating with vasectomized males. On the fourth day of pseudo-pregnancy, 20 μL of sesame oil was administered into one uterine horn, while the contralateral horn remained untreated, serving as a control. Eight days into pseudo-pregnancy, uterine samples were harvested for subsequent analysis.

### Quantitative RT-PCR

Total RNA was extracted utilizing the TRIzol reagent (Accurate Biotechnology). Genomic DNA was eliminated through treatment with DNase I (Invitrogen). The cDNA synthesis was achieved using the PrimeScript reverse transcriptase reagent kit (TaKaRa). Quantitative PCR reactions were performed on the Applied Biosystems 7500 platform (Life Technologies) with the THUNDERBIRD SYBR qPCR Mix (Toyobo). The Rpl7 gene served as the reference gene for normalization. The sequences of all primers used in this study are provided in **Table S3**.

### Western blot

The protein concentration was accurately measured using the BCA protein assay kit (Applygen). Subsequently, protein samples were separated via electrophoresis and efficiently transferred onto polyvinylidene difluoride (PVDF) membranes. To eliminate nonspecific binding, the membranes were blocked with 5% skim milk at room temperature for 1 hour. Subsequently, the membranes were incubated with the primary antibody overnight at 4°C. Following three washes with TBS-T containing 5% milk, the membranes were incubated with a horseradish peroxidase (HRP)-conjugated secondary antibody for 1 hour at room temperature. Chemiluminescent signals were detected using an ECL Chemiluminescent kit (Millipore). The intensities of the bands were precisely analyzed using the ImageJ software v1.54i [32]. Uncropped and unprocessed scans of the blots are presented in **Fig. S7**. The primary antibodies utilized in this study are comprehensively detailed in **Table S4**.

### Immunohistochemistry

Paraformaldehyde-fixed and paraffin-embedded tissues were delicately sliced into 5 μm sections. Antigen retrieval was meticulously conducted using citrate buffer adjusted to a pH of 6.0. To eliminate nonspecific binding, the sections were blocked with 10% horse serum diluted in PBS. Subsequently, the sections were incubated overnight at 4°C with the primary antibody to ensure optimal binding. After thorough rinsing with PBS three times to remove unbound antibody, the sections were incubated with a horseradish peroxidase (HRP)- conjugated secondary antibody for 1 hour at room temperature. The signal was then developed using the diaminobenzidine (DAB) kit (Zhongshan Golden Bridge Biotechnology Co., Beijing, China). Finally, the sections were counterstained with hematoxylin to enhance visualization. H-scores were accurately calculated using the QuPath software v0.5.1 [33]. The primary antibodies utilized in this study are comprehensively listed in **Table S4**.

### Alkaline phosphatase activity staining

To assess alkaline phosphatase activity, frozen tissue sections of 10 μm in thickness were utilized. The slides were fixed in cold paraformaldehyde (PFA) for 15 minutes. Subsequently, the sections were thoroughly washed three times with phosphate-buffered saline (PBS). For staining, a solution containing 5-bromo-4-chloro-3-indolyl phosphate (BCIP) and Nitro blue tetrazolium chloride (NBT), supplied by Zhongshan Golden Bridge Biotechnology Co., Beijing, China, was employed. After staining, the sections were counterstained with 1% methyl green to provide a clear contrast for visualization.

### RNA-seq

Total RNA was extracted using the TRIzol reagent (Invitrogen). The quality of extracted RNA was rigorously evaluated using both the ND-1000 Nanodrop and the Agilent 2200 Tape Station. RNA-seq libraries were meticulously prepared with the TruSeq RNA sample preparation kit (Illumina), following the manufacturer’s protocols. High-throughput sequencing was then performed on the HiSeq 2500 system (Illumina). After quality control, clean reads were mapped to the mouse genome (UCSC mm10) using Hisat2 v2.2.1 [34]. Mapped reads were subsequently analyzed with Cufflinks v2.2.1 [35] to reconstruct and quantify gene transcripts. Differentially expressed genes were identified using the following criteria: fold change > 2 and P < 0.05.

### Gene ontology analysis

The DAVID online tools [36] were utilized for gene ontology (GO) analysis. Genes were categorized based on their specific involvement in diverse biological processes. To ensure clarity and specificity, redundant GO terms were carefully removed through manual curation. A P-value cutoff of 0.05 was implemented to select significantly enriched GO terms.

### Statistical analysis

Statistical analysis was meticulously performed using GraphPad Prism v10.0.0 (https://www.graphpad.com/). To compare data between two groups, the Student’s t-test was utilized. For the analysis of multiple groups of data, a combination of one-way ANOVA and Tukey’s multiple comparison test was employed. To ensure reproducibility and consistency, all experiments were repeated at least three times. The data were presented as the mean ± SEM. Statistical significance was determined at a threshold of P < 0.05.

## Data availability

The sequencing data generated in this study have been deposited in the Gene Expression Omnibus (GEO) repository under the accession codes GSE263332.

## Author contributions

Y.-W.X., S.-H.Y. and J.-L.L. designed research; Y.-Q.H., R.-F.J., S.T., X.-Q.Y., M.Z., Y.L., and Q.-Y.Z. performed the experiments; Y.-Q.H., C.-H.D., Z.-H.Z., Y.-W.X., S.-H.Y. and J.-L.L. analyzed the data. Y.-Q.H., Y.-W.X., S.-H.Y. and J.-L.L. wrote the paper. All authors read and approved the final paper.

## Acknowledgments

This research was funded by National Natural Science Foundation of China (32370913 and 32070845), Guangdong Natural Science Funds for Distinguished Young Scholars (2021B1515020079), Innovation Team Project of Guangdong University (2019KCXTD001), Guangdong Special Support Program (2019BT02Y276), National Key R&D Program of China (2018YFA0801404), and Double first-class discipline promotion project (2023B10564003).

## References

1. Murakami K, Lee YH, Lucas ES, Chan YW, Durairaj RP, Takeda S, et al. Decidualization induces a secretome switch in perivascular niche cells of the human endometrium. Endocrinology. 2014;155(11):4542–53. Epub 2014/08/15. doi: 10.1210/en.2014-1370. PubMed PMID: 25116707.

2. Ramathal CY, Bagchi IC, Taylor RN, Bagchi MK. Endometrial decidualization: of mice and men. Semin Reprod Med. 2010;28(1):17–26. Epub 2010/01/28. doi: 10.1055/s-0029-1242989. PubMed PMID: 20104425; PubMed Central PMCID: PMCPMC3095443.

3. Wang H, Dey SK. Roadmap to embryo implantation: clues from mouse models. Nat Rev Genet. 2006;7(3):185–99. Epub 2006/02/18. doi: 10.1038/nrg1808. PubMed PMID: 16485018.

4. Gellersen B, Brosens JJ. Cyclic decidualization of the human endometrium in reproductive health and failure. Endocr Rev. 2014;35(6):851–905. Epub 2014/08/21. doi: 10.1210/er.2014-1045. PubMed PMID: 25141152.

5. Meyer KD, Jaffrey SR. The dynamic epitranscriptome: N6-methyladenosine and gene expression control. Nat Rev Mol Cell Biol. 2014;15(5):313–26. Epub 2014/04/10. doi: 10.1038/nrm3785. PubMed PMID: 24713629; PubMed Central PMCID: PMCPMC4393108.

6. Zaccara S, Ries RJ, Jaffrey SR. Reading, writing and erasing mRNA methylation. Nat Rev Mol Cell Biol. 2019;20(10):608–24. Epub 2019/09/15. doi: 10.1038/s41580-019-0168-5. PubMed PMID: 31520073.

7. Oerum S, Meynier V, Catala M, Tisne C. A comprehensive review of m6A/m6Am RNA methyltransferase structures. Nucleic Acids Res. 2021;49(13):7239–55. Epub 2021/05/24. doi: 10.1093/nar/gkab378. PubMed PMID: 34023900; PubMed Central PMCID: PMCPMC8287941.

8. Jia G, Fu Y, Zhao X, Dai Q, Zheng G, Yang Y, et al. N6-methyladenosine in nuclear RNA is a major substrate of the obesity-associated FTO. Nat Chem Biol. 2011;7(12):885–7. Epub 2011/10/18. doi: 10.1038/nchembio.687. PubMed PMID: 22002720; PubMed Central PMCID: PMCPMC3218240.

9. Zheng G, Dahl JA, Niu Y, Fedorcsak P, Huang CM, Li CJ, et al. ALKBH5 is a mammalian RNA demethylase that impacts RNA metabolism and mouse fertility. Mol Cell. 2013;49(1):18–29. Epub 2012/11/28. doi: 10.1016/j.molcel.2012.10.015. PubMed PMID: 23177736; PubMed Central PMCID: PMCPMC3646334.

10. Zheng ZH, Zhang GL, Jiang RF, Hong YQ, Zhang QY, He JP, et al. METTL3 is essential for normal progesterone signaling during embryo implantation via m(6)A-mediated translation control of progesterone receptor. Proc Natl Acad Sci U S A. 2023;120(5):e2214684120. Epub 2023/01/25. doi: 10.1073/pnas.2214684120. PubMed PMID: 36693099; PubMed Central PMCID: PMCPMC9945998.

11. Wan S, Sun Y, Zong J, Meng W, Yan J, Chen K, et al. METTL3-dependent m(6)A methylation facilitates uterine receptivity and female fertility via balancing estrogen and progesterone signaling. Cell Death Dis. 2023;14(6):349. Epub 2023/06/04. doi: 10.1038/s41419-023-05866-1. PubMed PMID: 37270544; PubMed Central PMCID: PMCPMC10239469.

12. Hong YQ, Tang S, Yang XQ, Deng YT, Zhao JQ, Zhang QY, et al. The METTL3–PGR– WNT4 axis is critical for human endometrial stromal cell decidualization. FEBS J. 2025. Accepted for publication 2025/04/11.

13. Wu Y, Su K, Zhang Y, Liang L, Wang F, Chen S, et al. A spatiotemporal transcriptomic atlas of mouse placentation. Cell Discov. 2024;10(1):110. Epub 2024/10/23. doi: 10.1038/s41421-024-00740-6. PubMed PMID: 39438452; PubMed Central PMCID: PMCPMC11496649.

14. Bachelot A, Binart N. Corpus luteum development: lessons from genetic models in mice. Curr Top Dev Biol. 2005;68:49–84. Epub 2005/08/30. doi: 10.1016/S0070-2153(05)68003-9. PubMed PMID: 16124996.

15. Soares MJ, Konno T, Alam SM. The prolactin family: effectors of pregnancy-dependent adaptations. Trends Endocrinol Metab. 2007;18(3):114–21. Epub 2007/02/28. doi: 10.1016/j.tem.2007.02.005. PubMed PMID: 17324580.

16. Lee KY, Jeong JW, Wang J, Ma L, Martin JF, Tsai SY, et al. Bmp2 is critical for the murine uterine decidual response. Mol Cell Biol. 2007;27(15):5468–78. Epub 2007/05/23. doi: 10.1128/MCB.00342-07. PubMed PMID: 17515606; PubMed Central PMCID: PMCPMC1952078.

17. Franco HL, Dai D, Lee KY, Rubel CA, Roop D, Boerboom D, et al. WNT4 is a key regulator of normal postnatal uterine development and progesterone signaling during embryo implantation and decidualization in the mouse. FASEB J. 2011;25(4):1176–87. Epub 2010/12/18. doi: 10.1096/fj.10-175349. PubMed PMID: 21163860; PubMed Central PMCID: PMCPMC3058697.

18. Yang Y, Zhu QY, Liu JL. Deciphering mouse uterine receptivity for embryo implantation at single-cell resolution. Cell Prolif. 2021;54(11):e13128. Epub 2021/09/25. doi: 10.1111/cpr.13128. PubMed PMID: 34558134; PubMed Central PMCID: PMCPMC8560620 as prejudicing the impartiality of the research reported.

19. He JP, Tian Q, Zhu QY, Liu JL. Identification of Intercellular Crosstalk between Decidual Cells and Niche Cells in Mice. Int J Mol Sci. 2021;22(14). Epub 2021/07/25. doi: 10.3390/ijms22147696. PubMed PMID: 34299317; PubMed Central PMCID: PMCPMC8306874.

20. Yang Y, He JP, Liu JL. Cell-Cell Communication at the Embryo Implantation Site of Mouse Uterus Revealed by Single-Cell Analysis. Int J Mol Sci. 2021;22(10). Epub 2021/06/03. doi: 10.3390/ijms22105177. PubMed PMID: 34068395; PubMed Central PMCID: PMCPMC8153605.

21. Jeong JW, Lee KY, Kwak I, White LD, Hilsenbeck SG, Lydon JP, et al. Identification of murine uterine genes regulated in a ligand-dependent manner by the progesterone receptor. Endocrinology. 2005;146(8):3490–505. Epub 2005/04/23. doi: 10.1210/en.2005-0016. PubMed PMID: 15845616.

22. Huyen DV, Bany BM. Evidence for a conserved function of heart and neural crest derivatives expressed transcript 2 in mouse and human decidualization. Reproduction. 2011;142(2):353–68. Epub 2011/04/30. doi: 10.1530/REP-11-0060. PubMed PMID: 21527398; PubMed Central PMCID: PMCPMC3141103.

23. Li Q, Kannan A, DeMayo FJ, Lydon JP, Cooke PS, Yamagishi H, et al. The antiproliferative action of progesterone in uterine epithelium is mediated by Hand2. Science. 2011;331(6019):912-6. Epub 2011/02/19. doi: 10.1126/science.1197454. PubMed PMID: 21330545; PubMed Central PMCID: PMCPMC3320855.

24. Wang C, Zhao M, Zhang WQ, Huang MY, Zhu C, He JP, et al. Comparative Analysis of Mouse Decidualization Models at the Molecular Level. Genes (Basel). 2020;11(8). Epub 2020/08/23. doi: 10.3390/genes11080935. PubMed PMID: 32823685; PubMed Central PMCID: PMCPMC7465532.

25. Liu JL, Wang TS. Systematic Analysis of the Molecular Mechanism Underlying Decidualization Using a Text Mining Approach. PLoS One. 2015;10(7):e0134585. Epub 2015/07/30. doi: 10.1371/journal.pone.0134585. PubMed PMID: 26222155; PubMed Central PMCID: PMCPMC4519252.

26. Murakami S, Jaffrey SR. Hidden codes in mRNA: Control of gene expression by m(6)A. Mol Cell. 2022;82(12):2236–51. Epub 2022/06/18. doi: 10.1016/j.molcel.2022.05.029. PubMed PMID: 35714585; PubMed Central PMCID: PMCPMC9216239.

27. Zhou Y, Zeng P, Li YH, Zhang Z, Cui Q. SRAMP: prediction of mammalian N6- methyladenosine (m6A) sites based on sequence-derived features. Nucleic Acids Res. 2016;44(10):e91. Epub 2016/02/21. doi: 10.1093/nar/gkw104. PubMed PMID: 26896799; PubMed Central PMCID: PMCPMC4889921.

28. Bastu E, Demiral I, Gunel T, Ulgen E, Gumusoglu E, Hosseini MK, et al. Potential Marker Pathways in the Endometrium That May Cause Recurrent Implantation Failure. Reprod Sci. 2019;26(7):879–90. Epub 2018/08/08. doi: 10.1177/1933719118792104. PubMed PMID: 30081718.

29. Xue P, Zhou W, Fan W, Jiang J, Kong C, Zhou W, et al. Increased METTL3-mediated m(6)A methylation inhibits embryo implantation by repressing HOXA10 expression in recurrent implantation failure. Reprod Biol Endocrinol. 2021;19(1):187. Epub 2021/12/16. doi: 10.1186/s12958-021-00872-4. PubMed PMID: 34906165; PubMed Central PMCID: PMCPMC8670269.

30. Koot YE, van Hooff SR, Boomsma CM, van Leenen D, Groot Koerkamp MJ, Goddijn M, et al. An endometrial gene expression signature accurately predicts recurrent implantation failure after IVF. Sci Rep. 2016;6:19411. Epub 2016/01/23. doi: 10.1038/srep19411. PubMed PMID: 26797113; PubMed Central PMCID: PMCPMC4726345 Organon, Schering Plough, Merck Serono, Ferring, Wyeth, Ardana, Andromed, Pantharei Bioscience and PregLem. NS Macklon has received fees and grant support from the following companies: Organon, Schering Plough, MSD, Anecova, IBSA, Merck Serono and Ferring. The other authors declare no competing financial interests.

31. Fukui Y, Hirota Y, Aikawa S, Sakashita A, Shimizu-Hirota R, Takeda N, et al. The EZH2- PRC2-H3K27me3 axis governs the endometrial cell cycle and differentiation for blastocyst invasion. Cell Death Dis. 2023;14(5):320. Epub 2023/05/18. doi: 10.1038/s41419-023-05832-x. PubMed PMID: 37198149; PubMed Central PMCID: PMCPMC10192223.

32. Schneider CA, Rasband WS, Eliceiri KW. NIH Image to ImageJ: 25 years of image analysis. Nat Methods. 2012;9(7):671–5. Epub 2012/08/30. doi: 10.1038/nmeth.2089. PubMed PMID: 22930834; PubMed Central PMCID: PMCPMC5554542.

33. Bankhead P, Loughrey MB, Fernandez JA, Dombrowski Y, McArt DG, Dunne PD, et al. QuPath: Open source software for digital pathology image analysis. Sci Rep. 2017;7(1):16878. Epub 2017/12/06. doi: 10.1038/s41598-017-17204-5. PubMed PMID: 29203879; PubMed Central PMCID: PMCPMC5715110.

34. Kim D, Paggi JM, Park C, Bennett C, Salzberg SL. Graph-based genome alignment and genotyping with HISAT2 and HISAT-genotype. Nat Biotechnol. 2019;37(8):907–15. Epub 2019/08/04. doi: 10.1038/s41587-019-0201-4. PubMed PMID: 31375807; PubMed Central PMCID: PMCPMC7605509.

35. Trapnell C, Williams BA, Pertea G, Mortazavi A, Kwan G, van Baren MJ, et al. Transcript assembly and quantification by RNA-Seq reveals unannotated transcripts and isoform switching during cell differentiation. Nat Biotechnol. 2010;28(5):511–5. Epub 2010/05/04. doi: 10.1038/nbt.1621. PubMed PMID: 20436464; PubMed Central PMCID: PMCPMC3146043.

36. Huang DW, Sherman BT, Tan Q, Kir J, Liu D, Bryant D, et al. DAVID Bioinformatics Resources: expanded annotation database and novel algorithms to better extract biology from large gene lists. Nucleic Acids Res. 2007;35(Web Server issue):W169-75. Epub 2007/06/20. doi: 10.1093/nar/gkm415. PubMed PMID: 17576678; PubMed Central PMCID: PMCPMC1933169.

