## Supplementary figures and tables for "Programmed downregulation of METTL3 is essential for decidualization in both humans and mice"

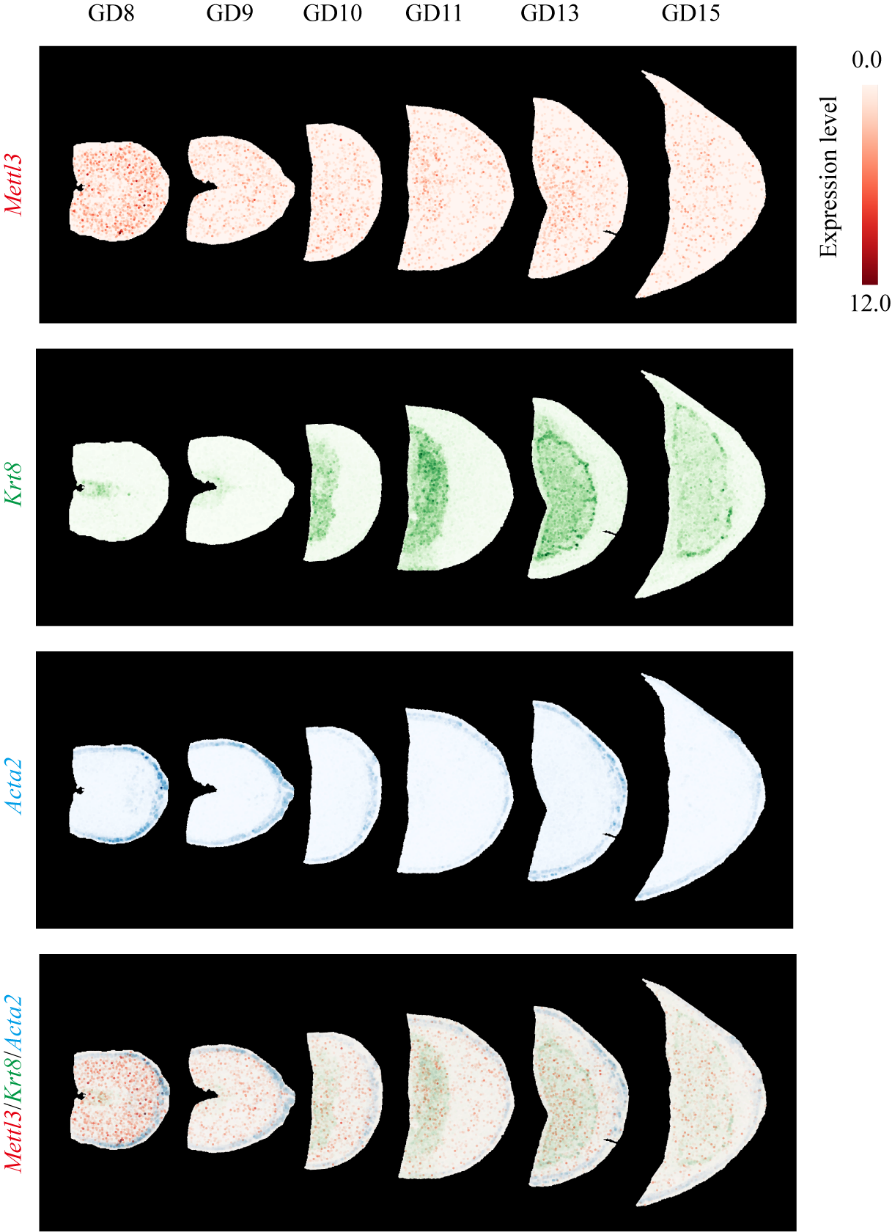

**Supplementary Figure 1. The expression pattern of *Mettl3* mRNA in the mesometrial decidua or decidua basalis across gestation days (GD) 8 to 15, as analyzed through a public Stereo-seq dataset.** The data is sourced from the Mouse Placentation Spatiotemporal Transcriptomic Atlas (https://db.cngb.org/stomics/mpsta/spatial/). GD1 denotes the day of vaginal plug detection. *Krt8* serves as a trophoblast cell marker for delineating the maternal-fetal interface, while *Acta2* marks smooth muscle.

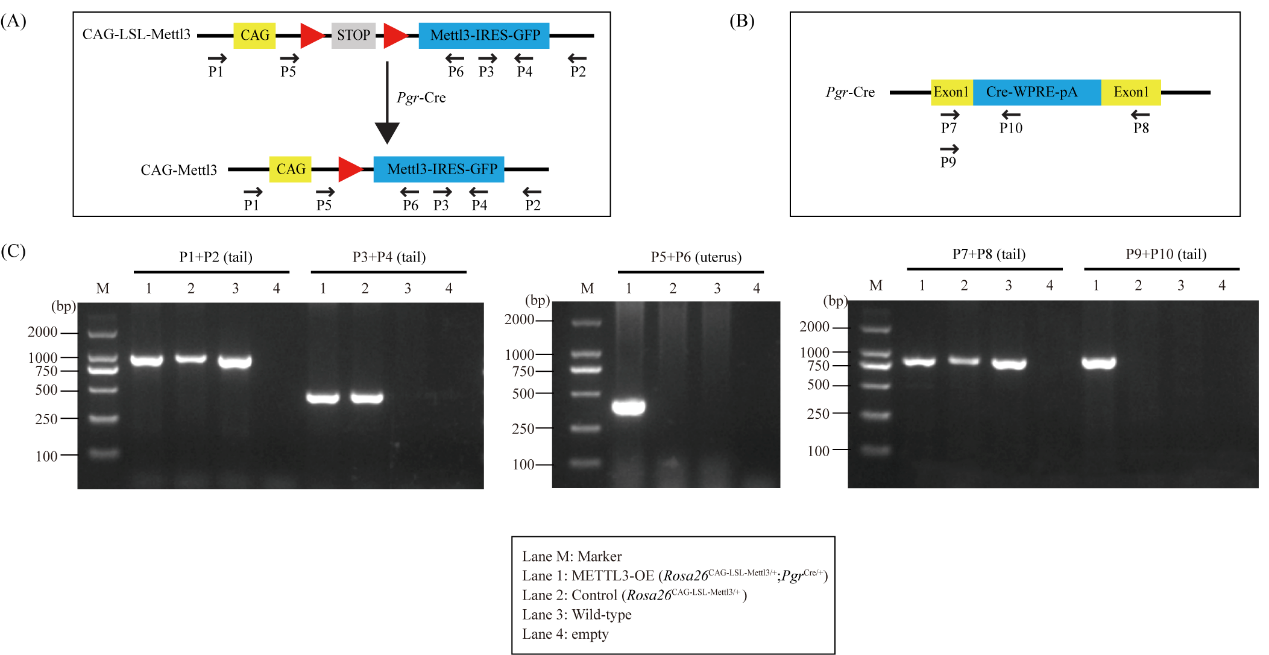

**Supplementary Figure 2. Genotyping analysis of METTL3-OE and control mice.** (A) Diagram showing genotyping primers for the METTL3 overexpression allele at the *Rosa26* locus. (B) Diagram showing genotyping primers for the *Pgr*-Cre allele. (C) PCR analysis for METTL3-OE and control mice.

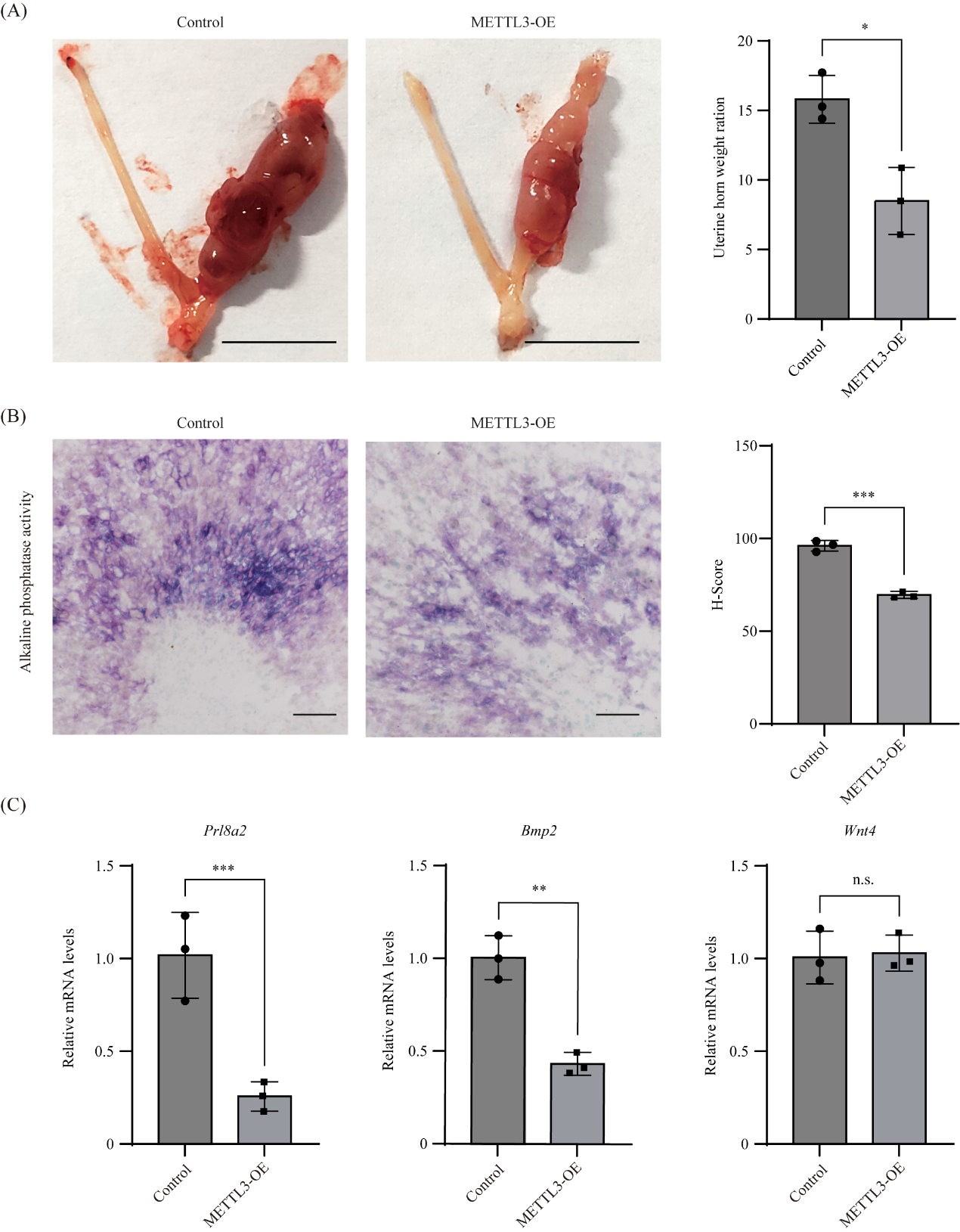

**Supplementary Figure 3. Compromised decidualization in METTL3-OE mice on GD8 can be recapitulated using an artificial decidualization model.** (A) Left panel: representative photos of uterine horns with or without stimulation from METTL3-OE and control mice. Bar = 1 cm. Right panel: bar plot showing the weight ratio between stimulated horn to the un-stimulated horn for METTL3-OE and control mice. Data are presented as mean ± SEM. *P < 0.05. (B) Alkaline phosphatase staining of the stimulated uterine horn from METTL3-OE and control mice. Bar = 100 μm. Data are presented as mean ± SEM. ***P < 0.001. (C) Quantitative RT-PCR analysis of *Prl8a2*, *Bmp2* and *Wnt4* in the stimulated uterine horn of METTL3-OE and control mice. Data are presented as mean ± SEM. **P < 0.01. ***P < 0.001. n.s., not significant.

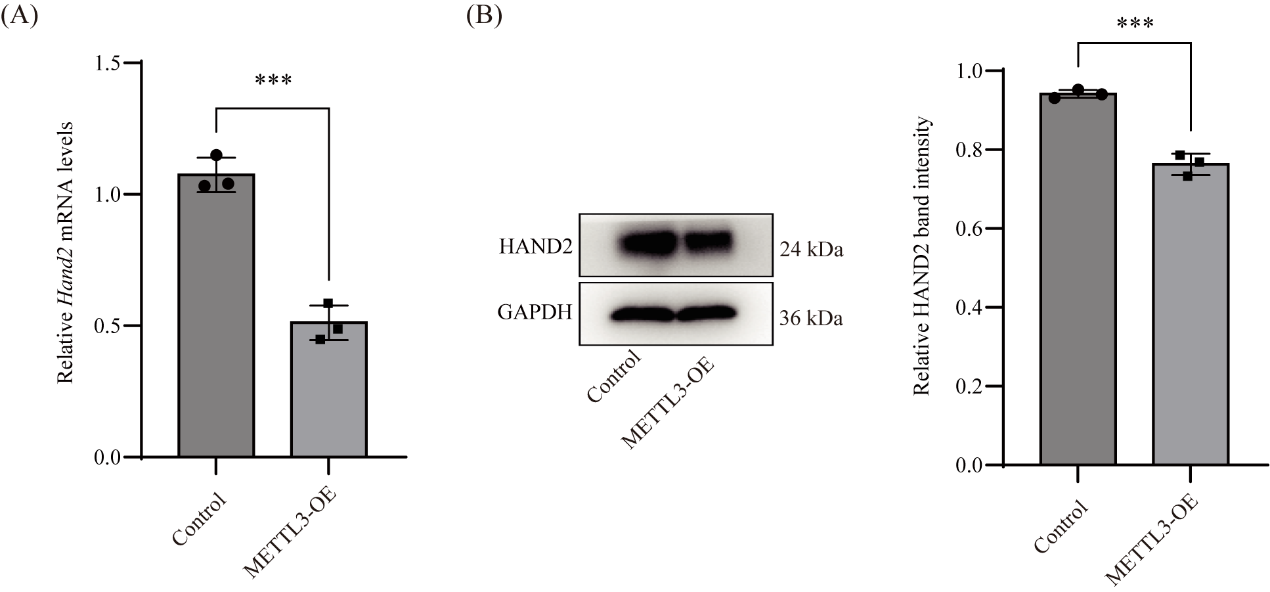

**Supplementary Figure 4. The expression of HAND2 in the uterus of METTL3-OE mice is decreased on GD4.** (A) Quantitative RT-PCR analysis of Hand2 mRNA levels in the uterus of METTL3-OE mice on GD4. Data are presented as mean ± SEM. ***P < 0.001. (B) Western blot analysis of HAND2 protein levels in the uterus of METTL3-OE mice on GD4. Data are presented as mean ± SEM. ***P < 0.001.

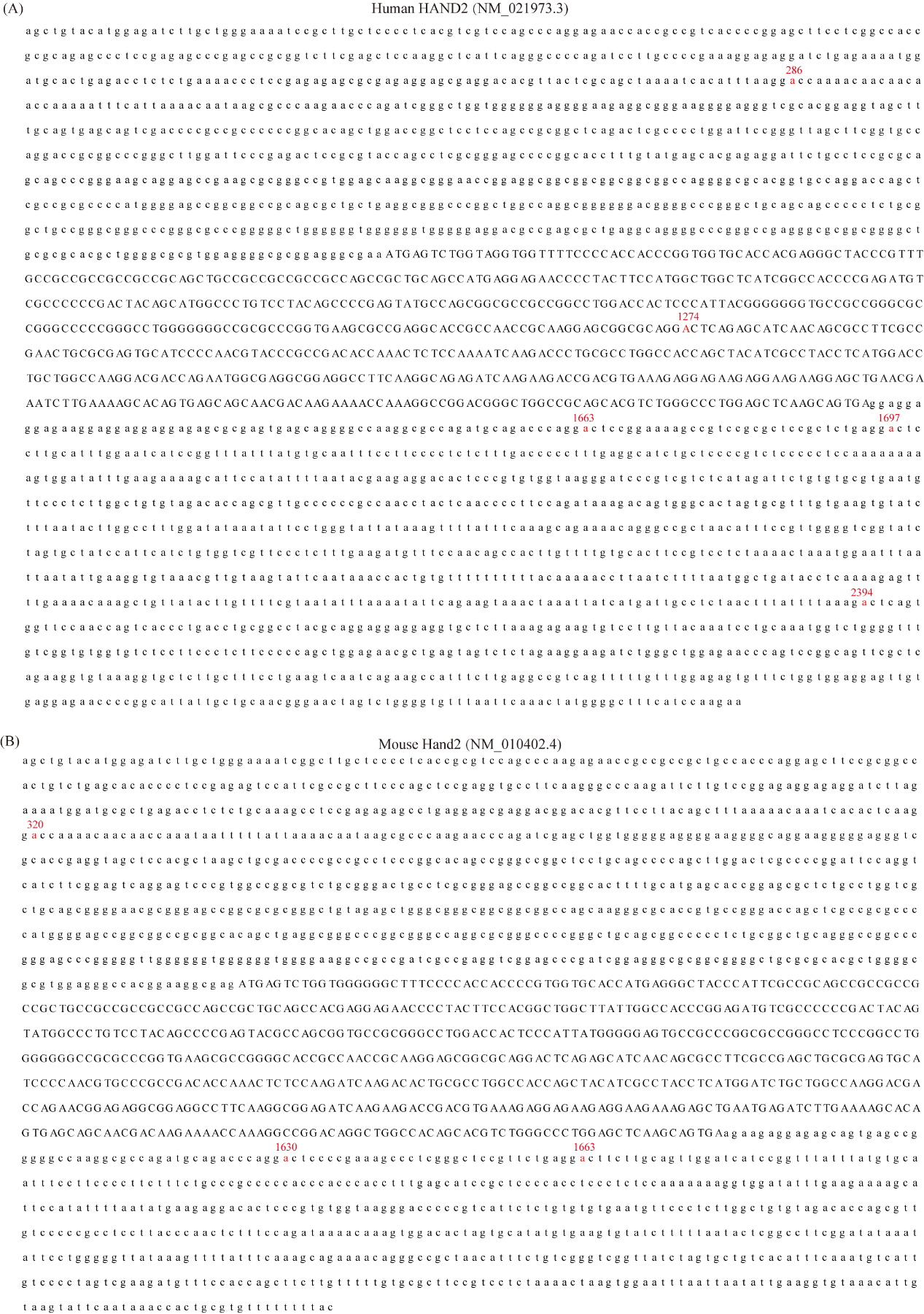

**Supplementary Figure 5. Identification of conserved potential m^6^A sites between humans and mice in HAND2 mRNA.** (A) The location of m^6^A sites in the 5′-UTR of human HAND2 mRNA predicted by the SRAMP tool (https://www.cuilab.cn/sramp). Potential m^6^A sites are colored in red. The 5′-UTR and the 3′-UTR is in lower case, while the coding sequence is in upper case. (B) Homologous potential m^6^A sites in mouse Hand2 mRNA identified by blast2seq (https://blast.ncbi.nlm.nih.gov/Blast.cgi). Human A286, A1663 and A1697 are homologous to mouse A320, A1630 and A1663, respectively.

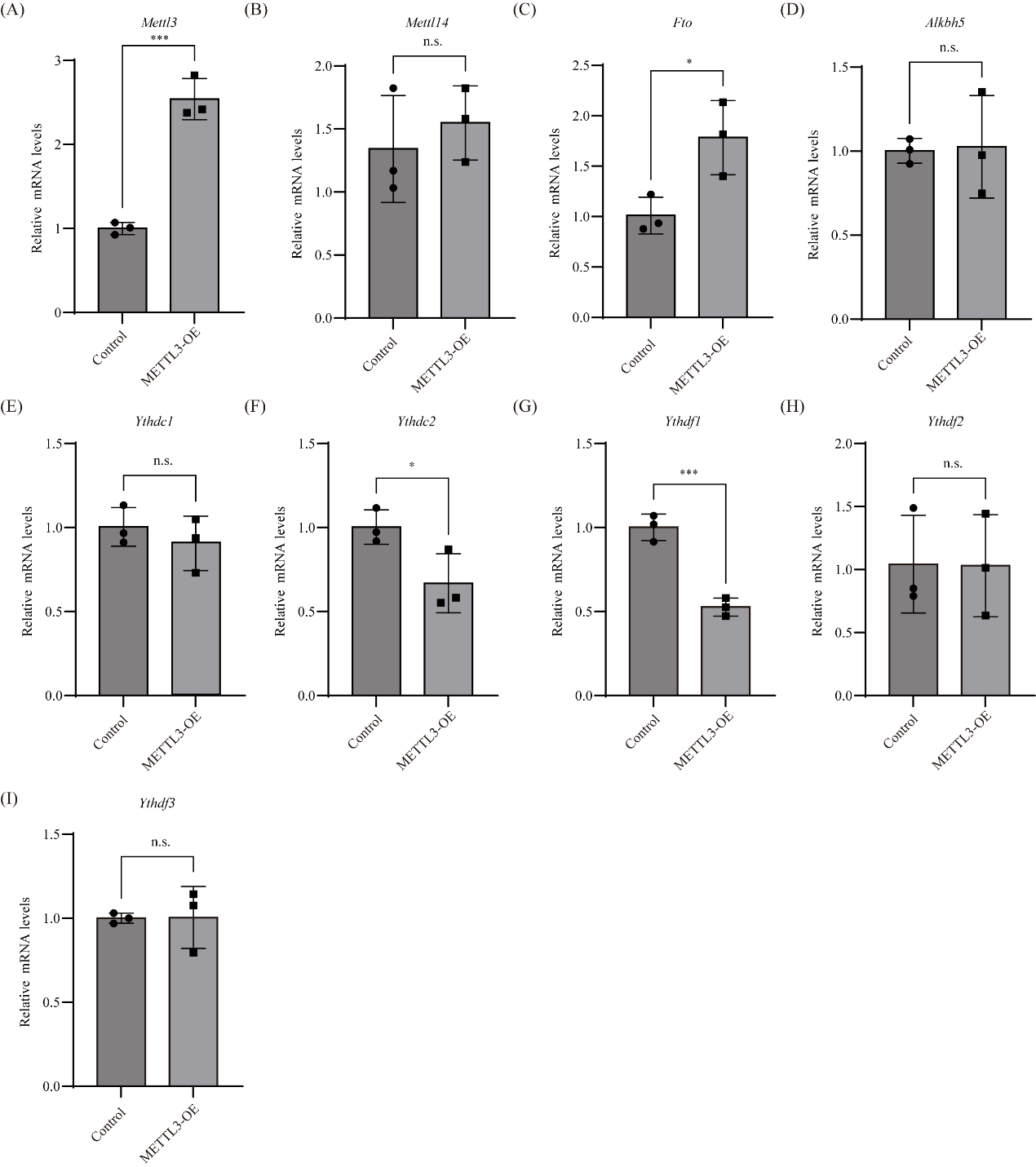

**Supplementary Figure 6. *Fto* is up-regulated, but *Ythdc2* and *Ythdf1* are down-regulated in the uterus of METTL3-OE mice on GD4.** Shown are qRT-PCR results for *Mettl3* (A), *Mettl14* (B), *Fto* (C), *Alkbh5* (D), *Ythdc1* (E), *Ythdc2* (F), *Ythdf1* (G), *Ythdf2* (H) and *Ythdf3* (I) in the uterus of METTL3-OE and control mice on GD4. Data are presented as mean ± SEM. *P < 0.05; ***P < 0.001.

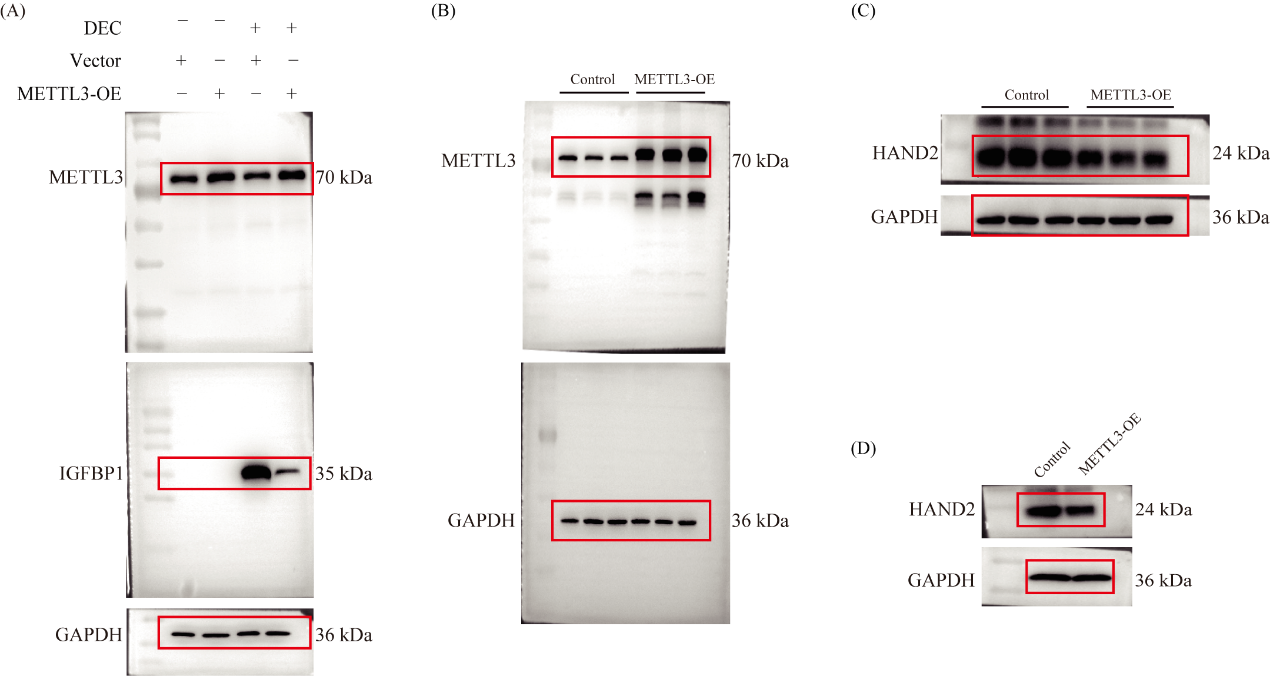

**Supplementary Figure 7. Images of the uncropped Western blots.** (A) Fig. 1B. (B) Fig. 2D. (C) Fig. 6E. (D) Fig. S4B

**Supplementary Tables**

**Supplementary Table 1. Differentially expressed genes in METTL3-OE uteri compared to control uteri on GD8 (fold change > 2 and P-value < 0.05).**

**Supplementary Table 2. Primers used in this study.**

**Supplementary Table 3. Antibodies used in this study.**

**Supplementary Table 4. Detailed information of human participants for isolation of primary endometrial stromal cells in this study.**
